# Intact memory for local and distal cues in male and female rats that lack adult neurogenesis

**DOI:** 10.1101/304733

**Authors:** Desiree R Seib, Erin Chahley, Oren Princz-Lebel, Jason S Snyder

## Abstract

The dentate gyrus is essential for remembering the fine details of experiences that comprise episodic memory. Dentate gyrus granule cells receive highly-processed sensory information and are hypothesized to perform a pattern separation function, whereby similar sensory inputs are transformed into orthogonal neural representations. Behaviorally, this is believed to enable distinct memory for highly interfering stimuli. Since the dentate gyrus is comprised of a large number of adult-born neurons, which have unique synaptic wiring and neurophysiological firing patterns, it has been proposed that neurogenesis may contribute to this process in unique ways. Some behavioral evidence exists to support this role, whereby neurogenesis-deficient rodents are impaired at discriminating the fine visuospatial details of experiences. However, the extent to which newborn neurons contribute to dentate gyrus-dependent learning tasks is unclear. Furthermore, since most studies of dentate gyrus function are conducted in male rats, little is known about how females perform in similar situations, and whether there might be sex differences in the function of adult neurogenesis. To address these issues, we examined spatial discrimination memory in transgenic male and female rats that lacked adult neurogenesis. The first task probed memory for the position of local objects in an open field, assessed by behavioral responses to novel object locations. The second task examined memory for distal environmental cues. All rats were able to successfully discriminate local and distal cue changes. Males and females also performed comparably, although females displayed higher levels of rearing and locomotion. Collectively, our results indicate that rats are capable of learning about local and distal cues in the absence of adult neurogenesis.

## INTRODUCTION

The dentate gyrus (DG) is the primary interface between the neocortex and the hippocampus proper. It receives highly-processed information about the external environment via the entorhinal cortex and is believed to perform a pattern separation function whereby similar inputs are transformed into distinct neural codes (Rolls and Kesner, 2006; Yassa and Stark, 2011). Behaviorally, this has been proposed to enable discrimination of similar sensory stimuli that are prone to interference.

While physiological assays have confirmed a pattern separation function for the DG, these experiments are technically challenging (Leutgeb et al., 2007; Neunuebel and Knierim, 2014). Thus, many have turned to behavioral tests of putative pattern separation-dependent, discrimination functions for the DG. In mice, deletion of DG NMDA receptors leads to impaired context discrimination in a fear conditioning paradigm (McHugh et al., 2007). DG-specific lesions and/or molecular manipulations also lead to impaired discrimination memory for object location and spatial geometry when stimuli are highly interfering (Gilbert et al., 2001; Hunsaker and Kesner, 2008; Bekinschtein et al., 2013). Such a discrimination function is not limited to animals, as imaging studies indicate that the DG-CA3 region is specifically recruited when subjects are asked to distinguish visual stimuli that are distinct, but bear a high resemblance, to previously-studied items (Bakker et al., 2008). Notably, in conditions such as aging and depression, which are associated with impaired hippocampal and DG function, there is poorer performance on human behavioral pattern separation tasks (Yassa et al., 2011; Déry et al., 2013; Shelton and Kirwan, 2013).

One feature that sets the DG apart from many regions of the mammalian brain is its ability to produce new neurons throughout adult life. These newborn neurons have enhanced synaptic plasticity in vitro, enhanced morphological plasticity in response to learning, and distinct wiring in hippocampal and entorhinal circuits (Vivar and van Praag, 2013). While the exact function for newborn neurons remains a topic of intense investigation, a number of animal studies suggest they may be involved in DG discrimination functions. Mice that have reduced or increased levels of neurogenesis show impaired or enhanced context discrimination abilities in fear conditioning paradigms, respectively (Sahay et al., 2011; Nakashiba et al., 2012; Tronel et al., 2012) (but see (Cushman et al., 2012; Snyder and Cameron, 2014)). Neurogenesis-deficient rodents are also impaired on spatial discrimination in radial maze and touchscreen paradigms (Clelland et al., 2009), and they are less able to learn lists of interfering odors (Luu et al., 2012). Finally, DG-specific manipulation of BDNF and Wnt signalling suggests that new neurons are required for remembering closely-related object locations (Bekinschtein et al., 2014).

While it remains unclear whether and/or how adult-born neurons might contribute to computational pattern separation (Aimone et al., 2011; Becker, 2017), these theoretical perspectives have played a significant role in guiding research on behavioral discrimination functions. The conditions under which new neurons contribute to behavioral discrimination remain unclear however. For example, many behavioral tasks vary in the degree of stress used for motivation, which could differentially recruit adult-born neurons (Snyder et al., 2011). Differences in behavioral requirements could also arise as a function of the testing paradigm and the sex of the animals. Indeed, in a context fear discrimination task, neurogenesis-deficient female mice discriminate normally but neurogenesis-deficient males show better discrimination learning than intact controls (Cushman et al., 2012). Also, in a radial maze, male rats that employed spatial strategies were better at discriminating similar locations and had greater neurogenesis compared to females (Chow, 2016). Here, we therefore tested male and female transgenic neurogenesis-deficient rats in a DG-dependent, object-location discrimination paradigm that measures memory purely based upon rodents’ natural tendency to explore novelty. In a second test, we examined rats’ ability to detect distal cue novelty. While our transgenic manipulation eliminated adult neurogenesis, we found that both male and female rats performed similarly, suggesting that adult neurogenesis is not critical for detecting at least some types of changes to the local and distal cue environment.

## MATERIALS AND METHODS

### Subjects

To examine behavioral functions of adult-born neurons we used GFAP-TK transgenic rats, in which neurogenesis can be selectively inhibited in adulthood via antiviral drug treatment (Snyder et al., 2016). Male and female GFAP-TK rats and their wild type littermates were bred in-house on a Long-Evans background, housed with their parents until they were weaned at 21 days of age, and kept on a 12-h light/dark cycle (lights on at 7:00 am). Experimental rats were pair housed in transparent polyurethane bins (48 × 27 × 20 cm) with a single polycarbonate tube, aspen chip bedding and ad libitum rat chow and tap water. The estrous cycle of female subjects was not monitored. To inhibit adult neurogenesis, at 6 weeks of age rats were orally administered 4 mg of valganciclovir (Hoffman La-Roche; delivered in 0.5 g peanut butter+chow pellets) twice per week. All experiments were approved by the Animal Care Committee at the University of British Columbia and conducted in accordance with the Canadian Council on Animal Care.

### Behavioral testing

Rats were treated with valganciclovir for 7-9 weeks (to inhibit DG neurogenesis) and began behavioral testing at 13 (males) or 15 (females) weeks of age (age difference because the sexes were tested separately; n=6-8/group; one male TK rat was excluded from the object location task because the power went out during testing).Valganciclovir treatment stopped once behavioral testing started, but neurogenesis remained reduced until the end of the study (Fig. 1). The week before testing started, animals were handled 5 min per day for 5 days. Rats were first tested in a proximal object-location discrimination test and then 7 days later a distal cue discrimination test (Fig. 1 D,E). All handling and behavior testing was conducted by female experimenters. On test days, animals were placed in the hallway in front of the experimental room 45 min before testing started. Both tests were performed in a square open field with transparent acrylic walls (70 cm wide × 70 cm long × 50 cm high), with aspen wood chip bedding covering the floor and 2 objects (sand-filled glass soda bottles) secured to the floor. The open field was placed in a dimly-lit small room (~2 × 2 m) with a door and posters on the walls to provide distal spatial cues. Both tasks consisted of 6 × 5 min trials. For the training trials (1-5) all environmental cues remained constant and rats were placed in an empty cage outside of the room for 3 min between each trial. There was a 10 min intertrial interval prior to trial 6, the testing trial, when cues were manipulated. The object-location discrimination task was modelled after Hunsaker et al. (2008) and Goodrich-Hunsaker et al. (2008). Objects were placed in the center of the open field and spaced 45 cm apart for trials 1 −5. On trial 6 the objects were shifted to 10 cm apart. In the distal cue novelty task the environment was identical to the object-location training trials (including bottle locations). However, on trial 6 dark curtains were hung on 2 distal room walls. In between trials objects were cleaned with 70 % EtOH, feces were removed, and bedding was stirred to prevent odors from accumulating. Between groups of male and females the open field was thoroughly cleaned and bedding was replaced.

**Figure 1:**
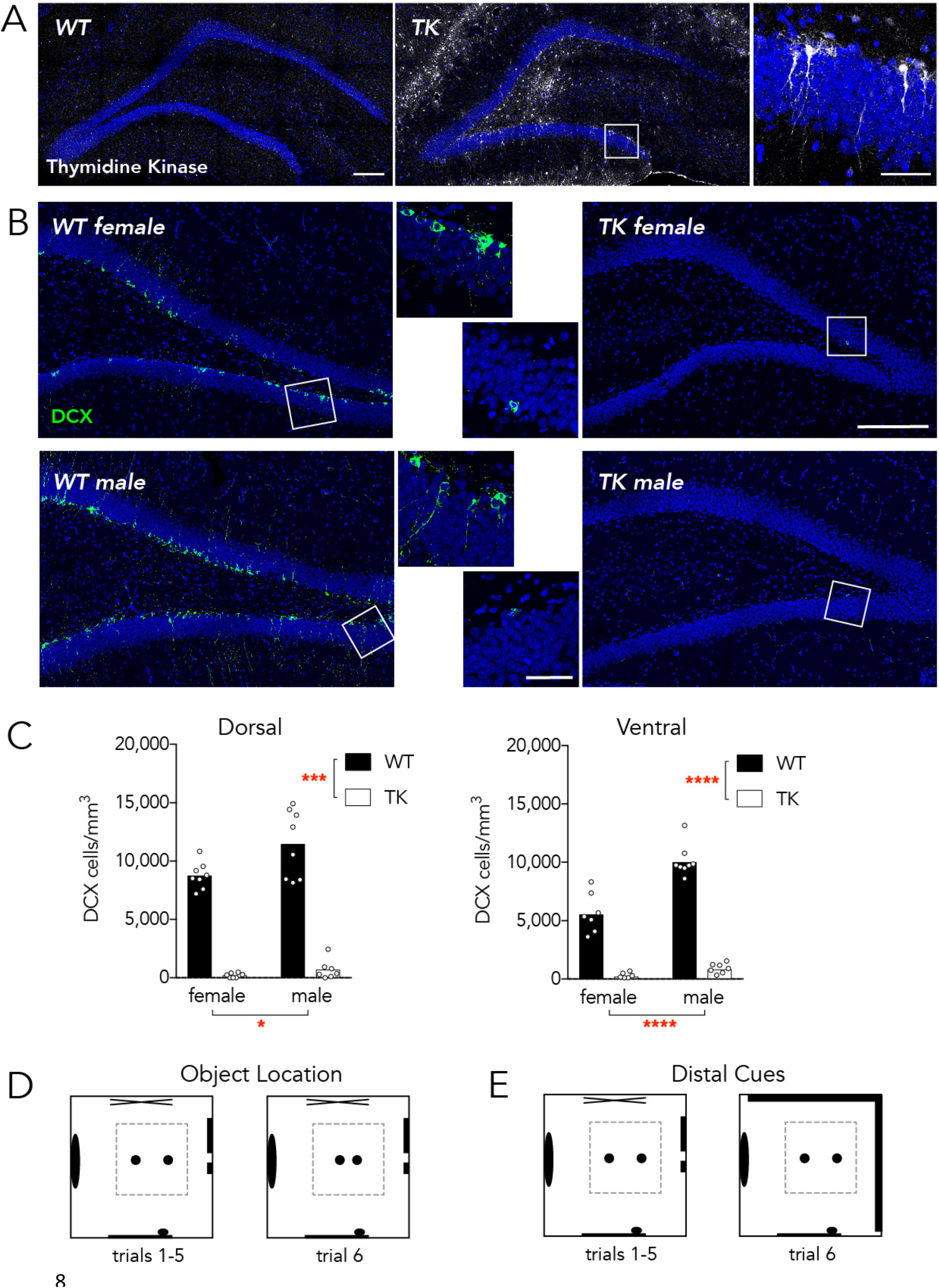
Neurogenesis knockdown and study design. A) Wild type and TK rats were immunostained for thymidine kinase after testing to confirm genotypes. Thymidine kinase-positive cells can be observed throughout the DG of TK rats. Inset shows thymidine kinase-positive radial glial cells in the subgranular zone. Scale bars: 200 μm for low magnification images, 50 μm for high magnification image. B) Male and female TK rats had dramatic reductions in neurogenesis, as visualized by immunostaining for the immature neuronal marker, DCX. Scale bars as in A. C) Mean DCX^+^ cell densities were reduced in TK animals and were greater in males than in females (Dorsal DG: effect of genotype F_1,27_=267, P<0.001; effect of sex F_1,27_=7.3, P<0.05; interaction F_1,27_=3.5, P=0.08; Ventral DG: effect of genotype F_1,27_ = 280, P<0.0001; effect of sex F_1,27_ = 34, P<0.0001; interaction F_1,27_=20, P=0.0002). In the ventral DG, all groups were different from each other (all P<0.0001) except TK males vs TK females (P=0.9) *P<0.05, ***P<0.001, ****P<0.0001. D) The object location discrimination test took place in a transparent-walled open field. During training, 2 bottles in the open field were spaced 45 cm apart during habituation trials 1 −5 and 10 cm apart during test trial 6. E) The distal cue discrimination test used the same habituation environment but for the test trial a dark curtain covered 2 of the room walls.

All trials were recorded by an overhead camera for offline analyses. The distance covered during each trial was measured by Ethovision software (Noldus). A human experimenter manually quantified the frequency of object exploration and rearing events. Exploration events were defined as continuous bouts of bottle sniffing and investigation (regardless of duration) where the rat’s nose was ≤ 2 cm from the bottle and the rat was directly exploring the bottle. Nondirected body/tail contact with the bottle, or contact when rearing (relatively rare) was not counted as bottle exploration. Rearing events, i.e. standing upright on hind legs, were interpreted as information-gathering behaviors directed at the distal environment (Lever et al., 2006). The absolute time spent investigating and rearing was also quantified but these data were more variable. Since the patterns and trends were closely correlated between the 2 measures we therefore focused our analyses on the frequency data. To further probe novelty responses on trial 6 compared to trial 5 we also calculated discrimination indices for each measure (trial 6/(trial 5+6)), which provided a single quantification of memory performance for each animal where values above 0.5 indicate greater-than-chance responses to novelty.

### Histology and microscopy

Following testing rat’s brains were extracted and immersed in 4 % paraformaldehyde for 48 hours. Brains were then transferred to a 10 % glycerol solution for 1 day and a 20 % glycerol solution for at least 2 days before being sectioned at 40 μm on a freezing sliding microtome. Dorsal hippocampal sections (~4 mm posterior to Bregma) were stained for thymidine kinase to confirm genotypes and dorsal and ventral (~6 mm posterior to Bregma) sections were stained for the immature neuronal marker doublecortin to confirm inhibition of neurogenesis (Fig. 1). Immunohistochemistry was performed with goat anti-HSV-1 thymidine kinase (sc-28038; Santa Cruz Biotechnology, USA) and goat anti-doublecortin (sc-8066; Santa Cruz Biotechnology, USA) primary antibodies, diluted in PBS with 0.05 % triton-x and 3 % normal horse serum, on free-floating sections. Secondary detection was performed with Alexa488-conjugated secondary antibodies (Life Technologies, USA) diluted in PBS. Sections were counterstained with DAPI, mounted onto slides, coverslipped with PVA-DABCO, and imaged with a Leica SP8 confocal microscope. DCX+ cell density measurements were calculated by dividing the number of DCX+ cells by the granule cell layer volume in one dorsal and one ventral section per animal.

### Statistical Analyses

Behavior across trials 1-6 was analyzed by 3-way repeated measures ANOVA. Mauchly’s test of sphericity was used to test for equal within-subject variance across trials and, when sphericity was violated, the Greenhouse-Geisser correction was applied. Where significant main effects or interactions were observed, post hoc comparisons were performed using Sidak’s test. Discrimination indices were analyzed by sex × genotype ANOVAs and individual groups were compared to chance using a one sample t-test. In all cases significance was set at p=0.05.

## RESULTS

### Neurogenesis reduction

Immunohistochemical staining for doublecortin (DCX) in the dentate gyrus following behavioral testing revealed extensive neurogenesis in valganciclovir-treated WT rats, as indicated by immature neurons scattered throughout the DG. In contrast, valganciclovir nearly completely inhibited adult neurogenesis in all male and female TK rats, in both the dorsal and ventral DG (Fig. 1 B,C). The density of DCX^+^ cells was greater in males than in females, in both the dorsal and ventral DG.

### Object Location discrimination task

For 5 consecutive trials, rats explored the open field with 2 objects spaced 45 cm apart; on the 6^th^ trial the distance between the objects was reduced to 10 cm. The first measure of performance in the object-location task was the frequency of object exploration events. There was a main effect of trial on object exploration but no effect of sex or genotype (see statistical analyses in figure legend). Rats explored the bottle less across the 5 habituation trials (Fig. 2A). Critically, exploration time doubled on trial 6 relative to trial 5, indicating that rats detected the new bottle locations. This pattern was also apparent in the discrimination indices, all of which were greater than chance, indicating successful discrimination of object location (Fig. 2B).

**Figure 2:**
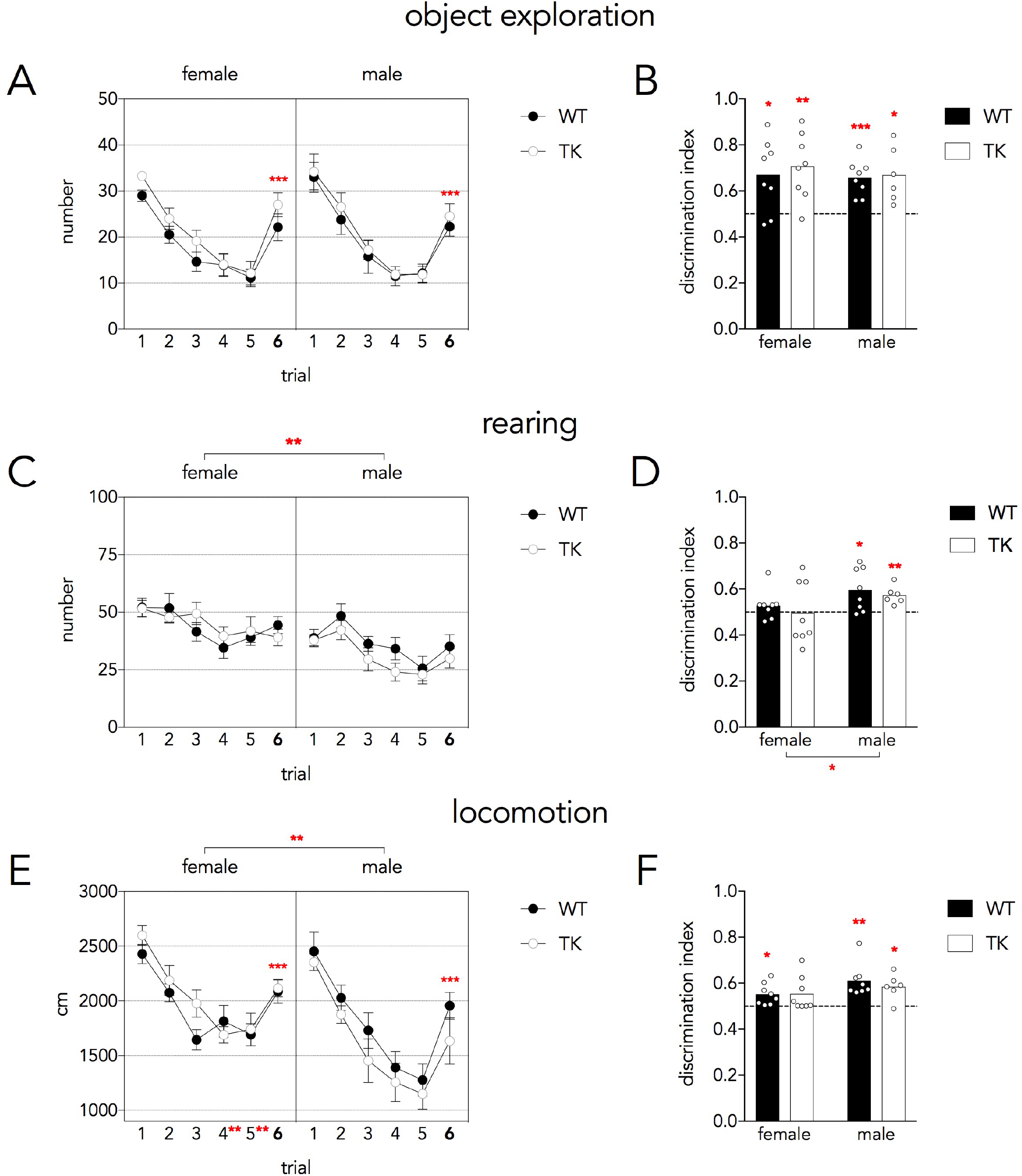
Object location discrimination task. A) The number of object exploration events decreased over the 5 habituation trials but doubled on trial 6 when object locations were changed (Mauchly s test P=0.3; effect of trial F_5,130_= 54, P<0.001, partial η^2^=0.68; trial 6 vs trials 3,4,5 all P<0.001). There were no differences between sexes, genotypes and no interactions (all P>0.16). B) All groups showed above-chance discrimination indices, indicating increased object exploration on trial 6 relative to trial 5. Variance was not different across groups (Levene’s test F_3,26_=1.9, P=0.16). C) The number of rearing events decreased over the 5 habituation trials (Mauchly’s test P=0.6; effect of trial F_5,130_=11, P<0.001, partial η^2^=0.30; trial 4&5 vs 1 &2 all P<0.001) but did not change between trials 5 and 6 (P=0.8). There were no genotype differences or interactions (all P>0.1) but females reared significantly more than males (F_1,26_=14, P<0.01, partial η^2^=0.35). D) The rearing discrimination index was above chance only in males and was greater in males than females (effect of sex F_1,26_=4. 5, P=0.04, partial η^2^=0.1 5). Due to unequal variance (Levene’s test P=0.002) and a lack of sex × genotype interaction, genotypes were pooled and the sex difference was reanalyzed, confirming a greater rearing discrimination index in males and homogeneity of variance across groups (T28=2.2, P=0.03; Levene’s P=0.25) E) Locomotion decreased over the 5 habituation trials in both sexes (Mauchly’s test P=0.4; effect of trial, F_5,130_ = 56, P<0.001, partial η^2^=0.68) and increased from trial 5 to trial 6 following object displacement (P<0.001). Locomotion was greater in females than males, specifically on trials 4 and 5 (effect of sex, F_1,26_= 10, P<0.01, partial η^2^=0.27; trial × sex interaction F_5,130_ = 2.3, P<0.05, partial η^2^=0.82; trials 4&5 different between males and females P<0.01). F) The locomotor discrimination index did not differ between genotypes (P=0.6, partial η^2^=0.01) or sexes (P=0.07, partial η^2^=0.12). Discrimination indices revealed above-chance levels of discrimination, except for female TK rats. Variance was not different across groups (Levene’s test F_3,26_=0.7, P=0.6). For all graphs: *P<0.05, **P<0.01, ***P<0.001. Graphs in panels A,C,E indicate mean ± standard error.

We next examined the frequency of rearing events. Again, we found a main effect of trial, indicating that rats habituated to the distal testing environment over the course of testing (Fig. 2C). Females reared 32% more than males, but the number of rearing events did not change between trials 5 and 6, and the rearing discrimination indices were at chance. In males, there was no significant increase in the absolute number of rearing events from trial 5 to 6. However, the discrimination indices were greater in males than in females (Fig. 2D). Moreover, only male discrimination indices were greater than chance, indicating sex differences in the behavioral response to local cue manipulations.

We then examined the distance travelled over the course of the 6 trials of the object-location task. There was a main effect of trial, with a significant increase in locomotion between trials 5-6 (22% in females, 48% in males), indicating that rats habituated to the environment and successfully detected the novel object location. While males and females initially covered similar distances on the early trials, males showed greater habituation over time such that locomotion was significantly reduced on trials 4 and 5 relative to females (Fig. 2E). Discrimination indices were not different between males and females or WT and TK rats, and most groups were significantly above chance performance (Fig. 2F).

### Distal cue novelty task

To test rats’ ability to detect changes in distal cues they were exposed to the open field for 5 consecutive training trials. On the 6^th^ trial a black curtain covered 2 adjacent walls (Fig. 1E). Over the 6 trials, rats explored the objects progressively less. They did not show any signs of increased bottle exploration following the distal cue manipulation, in terms of trial 5-6 differences or in the discrimination indices (Fig. 3A,B).

**Figure 3:**
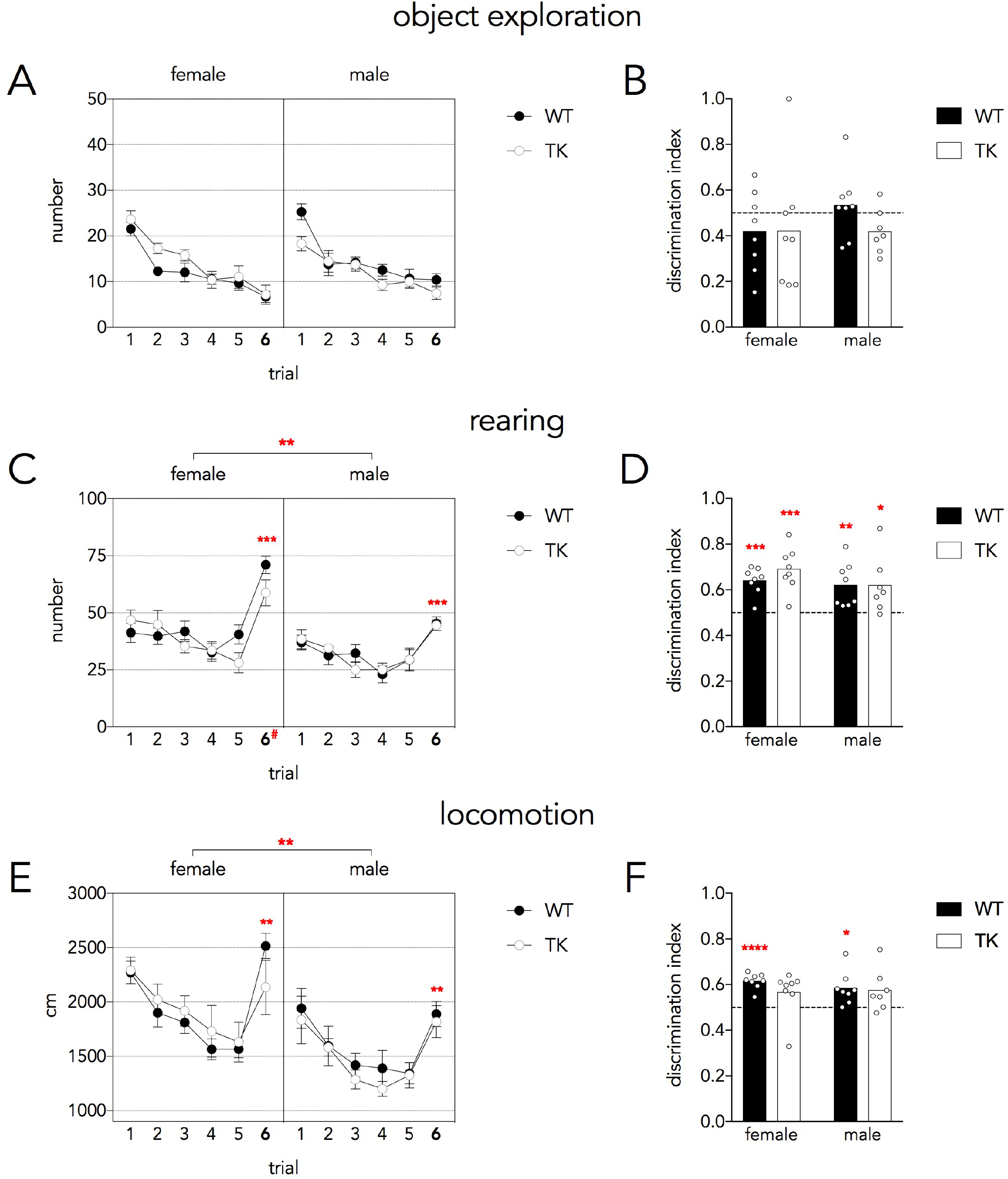
Distal cue novelty task. A) Rats decreased exploration of the objects over the 6 trials (Mauchly’s test P=0.2; trial effect, F_5,135_ = 44, P<0.001, partial η^2^=0.62) and did not increase exploration upon changes to the distal context on trial 6 (trial 6 vs trials 1 −4 all P<0.01, trial 6 vs trial 5 P=0.3). There were no genotype or sex differences (all P>0.5). B) Discrimination indices were not different between groups (genotype and sex effects P>0.4) and were not different from chance (all P>0.2), indicating that object exploratory behavior was unaffected by changes to distal cues. Variance was not different across groups (Levene’s test F_3,27_=1.5, P=0.25). C) Rearing events declined modestly over the 5 habituation trials and increased upon changes to the distal context on trial 6 (Mauchly’s test P=0.2; effect of trial, F_5,135_=32, P<0.001, partial η^2^=0.54; trial 6 vs all other trials P<0.001). Females reared more than males, particularly on trial 6 (sex effect F_1,27_=14, P<0.01, partial η^2^=0.33; trial × sex interaction F_5,135_=2.6, P=0.03, partial η^2^=0.087; females vs males on trial 6 #P<0.0001). WT and TK rats did not differ (F_1,27_=2, P=0.15, partial η^2^=0.015). D) Rearing discrimination indices did not differ between sexes or genotypes (both effects P>0.2) and all indices showed greater than chance levels of discrimination. Variance was not different across groups (Levene’s test F_3,27_=1.1, P=0.4). E) Distance travelled declined over trials until trial 6 when the distal cues were changed (Mauchly’s test P=0.001; Greenhouse-Geisser corrected trial effect F_2.9,135_=22, P<0.001, partial η^2^=0.45; trial 6 significantly greater than trials 3-5, P<0.01). Females covered a greater distance than males (sex effect F_1,27_=17, P<0.001, partial η^2^=0.39) and WT and TK rats did not differ (P=0.7). F) The locomotion discrimination index did not differ between genotypes or sexes (main effects P>0.3). Locomotion discrimination indices tended to be above chance levels of performance but only female TK and male WT indices were significantly above chance. Variance was not different across groups (Levene’s test F_3,27_=0.9, P=0.4). For all graphs: *P<0.05, **P<0.01, ***P<0.001, ****P<0.0001). Graphs in panels A,C,E indicate mean ± standard error.

The most robust response to the contextual change was observed in rearing behavior. Rearing decreased slightly over the first 5 trials but increased significantly on trial 6, indicating rats detected the changes to the distal context (Fig. 3C). Females reared significantly more than males (30% more overall), especially after the context change on trial 6. Rearing discrimination indices were above chance for all groups and did not differ between sexes or genotypes.

Locomotor behavior also changed in response to the distal cue changes. There was a decline in distance travelled over trials 1-5 but distance increased significantly on trial 6 (42% increase; Fig. 3E). Females covered 25% greater distance than males. Locomotor discrimination indices were consistent with distal cue novelty detection, though not all groups’ indices were statistically greater than chance (Fig. 3F).

## DISCUSSION

### Summary of main findings

Here we investigated the requirement of adult-born hippocampal neurons in an object location memory task that is designed to tap into DG functions in fine grained spatial processing. We also examined whether adult-born neurons are required to detect changes to the distal cue environment. Our principal finding is that adult neurogenesis was not required for learning either of these tasks. For the most part, males and females learned the tasks similarly. This is notable since no studies have systematically examined behavioral requirements for adult neurogenesis in male and females, and the vast majority of open field, novelty-based memory studies have exclusively used males. Consistent with previous studies, females were more active in the open field. Also, only males showed increased rearing behavior in response to local object location changes, suggesting sex differences in the behavioral response to local cue reconfigurations.

### Dentate gyrus and neurogenesis functions in memory

The DG is a major site of convergence of different forms of sensory information and, given the ability of DG neurons to undergo changes in synaptic strength, is well-positioned to mediate rapid encoding of sensory experience. Specifically, DG neurons receive inputs from the medial and lateral entorhinal cortices; the medial entorhinal cortex conveys spatial signals and information about self-motion and position in space (Fyhn et al., 2004; Hafting et al., 2005; Solstad et al., 2008). In contrast, the lateral entorhinal cortex conveys information about the identity and spatial location of objects in the environment (Deshmukh and Knierim, 2011; Tsao et al., 2013). The convergence of these inputs in the hippocampus produces high level representations that are the result of unique configurations of spatial, self and object-related cues (Eichenbaum et al., 2012; Hunsaker et al., 2013; Knierim et al., 2014). The specificity of DG place fields is consistent with this function and supports the computational role of the DG in pattern separation, whereby input patterns are orthogonalized to reduce memory interference (Neunuebel and Knierim, 2014). A critical behavioral role for the DG in spatial processing extends back to studies demonstrating that DG dysfunction leads to deficits in spatial water maze learning and context-specific expression of fear (Sutherland et al., 1983; Xavier et al., 1999; McHugh et al., 2007). A specific role for adult-generated DG neurons is supported by several studies that have found that they promote context-specific fear discrimination (Sahay et al., 2011; Kheirbek et al., 2012; Tronel et al., 2012), context-specific ensemble codes in CA3 (Niibori et al., 2012), memory for highly-interfering spatial locations (Clelland et al., 2009) (but see (Swan et al., 2014)) and lists of interfering odor pairs (Luu et al., 2012). Furthermore, emerging evidence suggests that mature neurons are more critical for behaviors that may rely on pattern completion functions whereas immature neurons necessary for behaviors that may rely more on pattern separation functions (Nakashiba et al., 2012; Danielson et al., 2016).

Here we exploited rats’ innate exploratory response to novelty (Ennaceur and Delacour, 1988). This approach has been used on numerous occasions to reveal functions for medial temporal lobe structures in memory for objects, contexts, time, and their combinations. An advantage of novelty-induced exploration is that it does not rely on aversive stimuli or food deprivation for motivation. This methodological difference is significant because adult-born neurons (and the hippocampus more broadly) regulate behavioral and endocrine responses to stressors (Snyder et al., 2011; Surget et al., 2011; Lehmann et al., 2013; Culig et al., 2017), and their functions in these more cognitive, less stressful, learning paradigms may therefore be distinct.

Novelty-induced exploration paradigms have identified roles for the DG and adult-born neurons in object-related memory. Rats with DG lesions, and mice with disrupted juvenile DG neurogenesis, are specifically impaired at detecting changes in local object positions when there is a high degree of spatial similarity between training and testing (Gilbert et al., 2001; Goodrich-Hunsaker et al., 2008; Hunsaker et al., 2008). Learning highly-interfering spatial object configurations upregulates BDNF specifically in the DG and both BDNF and new neurons have been implicated in 24 hour object location memory (Bekinschtein et al., 2013; 2014). Neurogenesis-deficient mice are impaired on (flexible) spatial discrimination of object-like cues presented on a touchscreen (Clelland et al., 2009; Swan et al., 2014). Notably, in addition to their role in remembering object location, adult-born neurons are also involved in object novelty detection (Jessberger et al., 2009; Denny et al., 2012; Bolz et al., 2015). Given the role of the lateral entorhinal cortex in these various types of object-related processing (Deshmukh and Knierim, 2011; Tsao et al., 2013), and its preferential connectivity with adult-born neurons (Vivar et al., 2012), there is broad support for a role for neurogenesis in object-related memory.

In our study, ablating adult neurogenesis did not impact rats’ ability to discriminate changes in the spatial positioning of objects. While this would seem to be at odds with previous studies, a number of methodological factors might have contributed to the differing reports. For example, deficits in the object-location discrimination task have been reported in mice that have had reduced neurogenesis since the juvenile period (Kesner et al., 2014). Behavioral effects may have been more apparent due to a greater net reduction in neurogenesis. Our task may have been easier than the one employed by Hunsaker et al. (2008) since we used 5 training trials instead of 3 (we found that rats did not learn reliably with only 3 training trials). Smaller spatial changes might also be more likely to reveal deficits, though ours were broadly comparable to those used previously and we did not even observe a trend for impaired performance in the TK rats. Our findings would seem to be at odds with the observation of Bekinschtein et al. (2014) that rats with reduced Wnt signalling and DG neurogenesis had impaired memory for object position. This could be due to differences in the spatial environment (they used a larger, circular open field with more objects and smaller changes to object location) or the testing schedule (theirs had only a single training trial and a longer, 24 hr, training-test interval) that could have made their task more difficult. Additionally, different methods to reduce neurogenesis could be at play (e.g. Wnt signalling contributes to synaptic plasticity (Ivanova et al., 2017)).

In contrast to the local cue manipulations of the first task, in the second task we manipulated the distal cue environment. In hippocampal-dependent tasks such as the water maze, distal cues are the primary source of visuospatial information.

Manipulation of the distal environment is also known to regulate hippocampal place fields (Leutgeb et al., 2005). Since, neurogenesis deficient animals have been found to be impaired in spatial tasks that rely on accurate memory for distal cues (Snyder et al., 2005; Clelland et al., 2009; Garthe et al., 2009), we reasoned that TK rats might show differences in responding to changes in the distal cue environment. However, as in the object discrimination task, neurogenesis-deficient rats performed normally. Given the role of the hippocampus in high-order, conjunctive encoding, future studies should expand the investigation to include novelty-based exploration tasks that require differential association of stimuli with spatial locations, contexts and temporal intervals (e.g. Langston and Wood, 2010).

### Behavioral patterns following local and distal cue changes

There were specific and complementary patterns of behavior depending on task and sex. Over the 5 learning trials in both tasks, all measures tended to decrease as rats habituated to the environment. Males and females tended to show similar habituation curves except for locomotion, where males habituated to a greater extent on the later trials (particularly in the object location task, where the open field was more novel). Females also displayed greater amounts of rearing and locomotion in both tasks. This is consistent with a large literature indicating that female rodents are more active than males. In open field environments, and when given access to running wheels, females consistently rear more and cover greater distance (Archer, 1975; Bartling et al., 2017; Basso and Morrell, 2017). As studies incorporating both males and females become more common, it will be important to account for activity differences when using novelty responses to assess memory (e.g. to ensure that high baseline activity does not obscure novelty responses).

The raw scores and discrimination indices revealed distinct behavioral patterns in the two tasks: object exploration increased only when object positions changed, rearing increased only when the distal cues changed, and locomotion increased following changes to both proximal objects and distal cues. The rearing discrimination index additionally revealed a male rearing response to the object location change. Since rearing is typically interpreted as a behavior that promotes investigation of the distal environment, this suggests that males may be more likely to re-explore the distal environment when there are changes to local cues. Considered another way, this finding may be consistent with evidence that women tend to have better object-related memory than men and, while rodent data are equivocal, it might also fit with reports of sex differences in object-location memory (reviewed in (Koss and Frick, 2017)).

## Conclusions

In sum, our results indicate that neurogenesis is not essential for short-term memory for local and distal cues, in males and females. These findings are in contrast with previous studies showing that the DG and neurogenesis contributes to various types of spatial discrimination memory. Functions of adult neurogenesis may be specific for certain types of conjunctive associations, and they could also be critically dependent on task parameters. Another possibility, is that new neurons may become involved in learning when there is a stressful component involved (Snyder et al., 2011). Given the fact that sex differences in hippocampal function emerge often in response to stress (Luine, 2002; Bangasser and Shors, 2007), it could be fruitful to investigate whether new neuron functions in (spatial discrimination) memory emerge in stressful situations.

